# Females outperform males in spatial learning despite increased amyloid plaques and microgliosis in a TgF344-AD rat model of Alzheimer’s disease

**DOI:** 10.1101/2022.03.27.485975

**Authors:** Osama Chaudry, Kelechi Ndukwe, Lei Xie, Peter A. Serrano, Maria E. Figueiredo-Pereira, Patricia Rockwell

## Abstract

Alzheimer’s disease (AD) is a progressive neurodegenerative disease and is the sixth leading cause of death in the US. AD is more prevalent in females than males. While estrogen provides neuroprotection in females, sex mediated differences in the development of AD pathology are not fully elucidated. Therefore, a comparison of the events that develop between sexes in the early-stage of AD pathology may reveal new potential targets for more effective therapeutic intervention. To address sex differences, we analyzed early stage 9-month male and female TgF344-AD (Tg-AD) rats, an AD model carrying the APPswe and Presenilin 1 (PS1ΔE9) mutations that develops progressive age-dependent AD pathology similar to humans. Using active place avoidance (aPAT) tests that assess hippocampal-dependent spatial learning and memory, we found significant deficits in Tg-AD females compared to wild type females, but no significant difference between the two male genotypes. Moreover, significant sex differences were observed in that Tg-AD females outperformed Tg-AD males in several measures of the aPAT test. Unexpectedly, Tg-AD females displayed higher levels of hippocampal amyloid plaques and amoeboid microglia than their Tg-AD male littermates. Furthermore, Tg-AD females experienced less hippocampal neuronal loss and had higher GluA2 subunit levels than Tg-AD males. Based on our findings, we propose that estrogen may protect females against cognitive impairment at early stages of AD by regulating GluA2 levels independently of amyloid plaque deposition and gliosis. Elucidating this potential protective mechanism of action of estrogen in AD could lead to new targets for early intervention.

## Introduction

Alzheimer’s disease (AD) is a neurodegenerative disease that is the major cause of dementia in the US. Historically, it is considered that women are at greater risk of developing AD than men as two out of three AD patients in the US are females [1]. One of the factors contributing to this sex-dependent disparity is that women have a longer life expectancy than men, and as aging is a major risk factor for AD it is not surprising that the risk of AD is greater for women [35]. Besides the aging factor, other sex differences in AD are supported by clinical, neuroimaging and pathological studies [21]. To improve disease management and potential treatments, there is a critical need to understand mechanisms and pathways that contribute to protection or to the greater risk for women to develop AD.

AD is characterized by the following hallmarks: cognitive deficits, extracellular amyloid beta (Aβ) plaques, intracellular neurofibrillary tangles, neuroinflammation, and neurodegeneration. In the current study we focus on all of these except neurofibrillary tangles, as they appear later in the progression of the disease. AD pathology starts decades before the onset of clinical symptoms, and the entorhinal cortex and hippocampus are among the initial and most extensively affected brain regions [48]. The course of the disease from early to late stages often follows the classical trisynaptic pathway from entorhinal cortex (layer II) → dentate gyrus → CA3 → CA1 [30]. This trysynaptic pathway plays a role in new memory acquisition and is highly vulnerable to premature degeneration [30]. The hippocampus is a region of the brain that possesses high levels of plasticity, which correlates with its extreme vulnerability to insult including head trauma, ischemia, seizures and severe stress [33]. These two aspects of hippocampal function, i.e. plasticity and vulnerability, are related to circulating glucocorticoids acting through their highly abundant hippocampal receptors, and excitatory amino acids acting as neurotransmitters [33]. Elucidating the relationship between hippocampal plasticity/vulnerability and AD pathology in the early stages of the disease could lead to new targets for early intervention.

Neurodegeneration and neuronal loss are prominent AD pathological features and are associated with the formation of amyloid plaques involving aggregation of Aβ peptides [13]. The accumulation of Aβ peptides is a result of dysregulated proteolytic processing of the amyloid precursor protein (APP) [32]. APP is a transmembrane protein and although its exact physiological function remains speculative, it is implicated in neurite outgrowth, synaptogenesis, axonal protein trafficking, transmembrane signal transduction, cell adhesion, and calcium metabolism [32, 55, 56]. APP cleavage by α-secretase within the Aβ domain at the Lys16-Leu17 bond produces sAPPα and an αCTF fragment which is further cleaved by γ-secretase to produce p83, impeding production of amyloidogenic Aβ peptides [55]. In AD, APP processing is associated with cleavage by BACE1, a major β-secretase [50]. Cleavage by BACE1 produces a βCTF fragment and sAPPβ, followed by γ-secretase cleavage of the βCTF fragment to produce Aβ, yielding Aβ-40 and Aβ-42. The latter peptide is hydrophobic and highly prone to aggregation due to the isoleucine and alanine amino acids at its C-terminus, distinguishing Aβ-42 from Aβ-40 [55]. Although AD is a multifactorial disease, Aβ dyshomeostasis is proposed to be one of the major triggers of the cascade of neurotoxic events leading to neurodegeneration in AD [45].

Apart from senile plaques, AD is linked to neuroinflammation as observed in post-mortem brain tissue of AD patients [19]. Inflammation is an innate mechanism activated in response to infection, toxins, and injury. In AD, however, there is an imbalance of pro-inflammatory and anti-inflammatory signaling, in part due to microglia-released cytokines [10], allowing inflammation to go unchecked and to become chronic [42]. Under steady-state conditions, microglia remain in a surveillance state, and have small soma and long processes [18]. Upon activation by a threat to the CNS, there is a microglia morphological change, triggering the release of pro-inflammatory cytokines [11]. In AD, it is hypothesized that Aβ leads to the activation of microglia, which localize to plaques to phagocytose them [5]. However, after long periods of time, microglia become enlarged and unable to clear the plaques, leading to prolonged inflammation, neuronal damage, and exacerbated AD pathology [24, 34, 46].

To address potential sex differences in the early stages of AD pathology, we chose 9-month male and female TgF344-AD (Tg-AD) rats. This transgenic rat model of AD develops amyloid plaques and gliosis in the hippocampus and cerebral cortex at 6 months of age, and neurofibrillary tangles, neuronal loss and learning deficits around 15-months of age [7]. The Tg-AD rat model was generated on a Fischer 344 background and expresses two human genes driven by the mouse prion promoter, APPswe (Swedish) mutation, and human presenilin-1 exon 9 deletion (PS1ΔE9) [7, 14]. Tg-AD rats express 2.6 times more human APP and 6.2 times more human PSEN1 in the rat brain than their endogenous rat counterparts [7]. The APPswe mutation promotes abnormal cleavage of APP by BACE1, increasing intraneuronal Aβ-42 deposition and soluble Aβ oligomers and plaques in the hippocampus and cortex of Tg-AD rats [7, 47]. The PS1ΔE9 mutation shifts Aβ generation to longer and more aggregation-prone Aβ peptides via changes to the γ-secretase complex [53]. The TgF344-AD rat model is unique as it develops the full array of AD pathology in a progressive and age-dependent manner, thus mimicking disease progression in humans [7].

We performed cognitive assessment on the 9-month Tg-AD rats in addition to immunohistochemical and western blot analyses of hippocampal tissue in these animals, to determine amyloid plaque load, microgliosis and neuronal loss, along with levels of GluA2 subunit of the AMPA receptor and the neuronal postsynaptic scaffolding protein PSD95. Both GluA2 and PSD95 levels were examined since GluA2 is increased by estrogen and is associated with improved spatial memory while PSD95 plays an important role in maintaining synapse structure and function. Our findings support the notion that in the early stages of AD, estrogen in females is protective against cognitive impairment, but has less of an effect on other pathological deficits linked to AD. Elucidating this potential protective mechanism of action of estrogen in AD could lead to new targets for early intervention.

## Materials and Methods

### TgF-344AD transgenic rat model of AD

Fisher transgenic F344-AD (Tg-AD) rats [7] expressing human Swedish amyloid precursor protein (APPswe) and Δ exon 9 presenelin-1 (PS1ΔE9) along with wild-type (WT) littermates were used for this study. The Tg-AD rats and wild type (WT) littermates were purchased from the Rat Resource and Research Center (RRRC, Columbia, MO), and arrived at Hunter College when they were approximately 4 weeks of age. The rats were housed in pairs on a 12h light/dark cycle with food and water available *ad libitum* and maintained at the Hunter College Animal Facility. All experiments were performed in compliance with the regulations of the Institutional Animal Care and Use Committee at Hunter College.

### Experimental Design

Male and female rats at the age of 9-months were studied in the following numbers: Tg-AD males n = 10, WT males n = 5, Tg-AD females n = 6, WT females n = 5. Hippocampal-dependent cognitive deficits were assessed with the active-place avoidance task (aPAT). Following behavioral testing, the rats were sacrificed, the brains rapidly isolated and bisected into hemispheres, and processed for the different assays as described below.

### Active Place Avoidance Task (aPAT)

This hippocampal-dependent cognitive test is an active test that uses negative reinforcement (electrical shock) to access spatial learning and memory performance [28]. aPAT assesses reference memory because long-term spatial learning is used as the rats experience repeated trials with a fixed shock quadrant, so referencing previous trials will aid in better performance. A computer-controlled system was used for aPAT (Bio-Signal Group, Acton, MA).

This test uses a dry circular arena revolving at a speed of one revolution/min with four quadrants, one of which is a shock zone, giving an electric shock of 0.2 amps upon animal entrance. The arena was enclosed with a fixed transparent plastic wall allowing the rats to navigate using spatial cues in the test room. An overhead camera (Tracker, Bio-Signal Group) was calibrated to the white hue of the rats and tracked the rats movement. Rats were given a 10-min habituation trial without a shock, allowing them to explore the platform and get accustomed to the visual cues. A 10-min break followed the habituation and then six training trials were performed (alternating 10-min in the arena and 10-min resting off the arena) with the shock on. Latency to first entrance, latency to second entrance, maximum time to avoid shock, number of entrances in shock zone, and number of shocks were recorded during the six training trials and during the test trial 24-hours later. The system software recorded data for all trials, and all data was exported to .tbl files and analyzed offline (TrackAnalysis, Bio-Signal Group).

### Brain tissue preparation and immunohistochemical analysis (IHC)

At 9-months of age, the rats were anesthetized (i.p.) with ketamine (100 mg/kg) and xylazine (5 mg/kg) and transcardially perfused for 15-min with cold RNAse-free PBS. The rat brains were removed, and the left hemisphere was micro-dissected (regions: prefrontal cortex, cingulate cortex, entorhinal cortex, and hippocampus) and immediately snap frozen for molecular or biochemical analyses. The right hemispheres were sequentially post-fixed with 4% paraformaldehyde for 48-hours at 4°C, cryoprotected in a 30% sucrose/PBS solution at 4°C until they sank to the bottom of the vial, flash frozen in 2-methylbutane, and stored at -80°C until sectioned. The right hemispheres were sectioned with a Leica CM 3050S cryostat. Hippocampal coronal sections of 30 μm in thickness were collected serially along the anteroposterior axis and stored at -20°C in cryoprotectant [30% glycerol (Fisher BioReagents, cat# 15514029) and ethylene glycol (Fisher BioReagents, cat# 10532595)] in PBS until use.

IHC was restricted to dorsal hippocampal tissue within the following Bregma coordinates: -3.36 mm to - 4.36mm [38]. Sections were processed with a mounted protocol for IHC analyses as described in [3]. Briefly, hippocampal sections were washed in 1X PBS for 5-min three times, followed by mounting and quenching with 0.05M glycine (Fisher BioReagents, cat# BP3815) solution for 30-min. Sections were washed with 1X PBS/Triton X-100 0.3% (Life Technologies, Thermo Fisher Sci., cat# PI85112) three times for 5-min and then blocked for 30-min with 30% normal goat serum (Vector Labs, cat# S-1000) and 1X PBS/Triton 0.3%. Two to three sections (averaged) from each rat were immunostained for Aβ plaques, microglia (Iba1 antibody), mature neurons (NeuN antibody), GluA2 subunit, and PSD95. Following the overnight incubation with the primary antibodies, sections were washed three times for 5-min with 1X PBS/Triton 0.03% and blocked for 15-min in 30% NGS and 1X PBS/Triton 0.3%. After blocking, sections were incubated for 60-min in the fluorescent secondary antibodies. Primary and secondary antibodies are listed in Supplemental Table 2. All antibodies were diluted in 30% NGS and 1X PBS/Triton 0.03%. Two sets of three washes at 5-min each were performed after the secondary antibody incubation, with 1X PBS/Triton 0.03% and 1X PBS, respectively. Mounting media with DAPI (VectaShield, cat# sku H-1200-10) was used to mount, and sections were stored in the dark at 4°C.

### Image Processing and Quantification

Sections were viewed on a Zeiss Axio Imager M2 microscope with AxioVision 4 module MosaiX software to capture ZVI files of 10x mosaic images of the whole hippocampus using a Zeiss AxioCam MRm Rev. 3 camera connected to a motorized stage. Signal density (O.D.) was quantified using Image J as previously described [8]. Each channel was analyzed to an antibody specific threshold. GFP spectrum filter set was used for Iba1 imaging, DSRed set was used Aβ imaging, and DAPI set was used for DAPI imaging. Exposure time for each channel was kept consistent between sections. For each captured image, ZVI files were loaded onto FIJI (Fiji Is Just ImageJ, NIH, Bethesda, MD) and converted to .tiff files for analyses. Images were analyzed to extract the positive signal from each image with custom batch-processing macroscripts created for each channel/marker using the following formulae: average pixel intensity + [(1.5 *[Iba1]*, 4.0 *[Aβ], or (1[GluA2, PSD95, NeuN]*) x Standard deviation of intensities]. FIJI-ImageJ was used for all image adjustment and for quantification of microglia and plaque size, count, and percentage area in the cornu ammonis 1 (CA1), cornu ammonis 3 (CA3), dentate gyrus (DG) and subiculum (SB) hippocampal regions.

### Microglia Analysis

Microglia exhibit a variety of morphologies that can be associated with their functions and distributed into three different groups according to their form factor (FF) which is defined as 4π X area/perimeter^2^ [9]. Each of the three microglia groups is defined as follows: *Ramified*, FF: 0 to 0.49; which actively engage in neuronal maintenance providing neurotrophic factors. *Reactive*, FF: 0.50 to 0.69; which are responsive to CNS injury, and *amoeboid*, FF 0.70 to 1; amorphous with pseudopodia. Microglia within each cropped Iba1 image were extracted using the following formula: average pixel intensity + [1.5 x standard deviation of intensities], and particles within 50– 800 µm^2^ were chosen for FF analyses. Nonspecific background density was corrected using ImageJ rolling-ball method [49].

### Western blot analysis

Hippocampal tissue (20-25 mg) was homogenized in TBS for 90 sec at 25°C with the Bedbug Microtube Homogenizer (3,400 rpm, Model D1030, Benchmark Scientific). The supernatant was stored for 16 h at -80°C, followed by centrifugation at 14,000 rpm for 20 min at 4°C. The supernatant was filtered using biomasher homogenizer tubes (#09-A10-050, OMNI International). Samples were stored at -80°C until use. Protein concentration was determined with the BCA assay (Pierce Biotechnology), followed by normalization. 30 μg from each sample were run on 4-12% SDS gels and transferred to nitrocellulose membranes with the iBlot^®^ dry blotting system (Life Technologies) for 7 min. Membranes were blocked with SuperBlock (#37535, ThermoFisher), and hybridized with various primary antibodies followed by HRP-conjugated secondary antibodies (Supplemental Table 2), prior to developing with an enhanced chemiluminescence (ECL) substrate (SuperSignal(tm) West Pico PLUS, ThermoFisher #34580), and detected on a BX810 autoradiography film (Midwest Scientific). ImageJ software (Rasband, W.S., ImageJ, U. S. National Institutes of Health, Bethesda, Maryland, USA, https://imagej.nih.gov/ij/, 1997-2018) was used for semi-quantification by densitometry of the respective bands. Loading controls used were GAPDH or β-actin depending on their molecular weights to avoid overlapping with the other proteins studied.

### Statistics

All data are represented as the mean ± SEM. Statistical analyses were performed with GraphPad Prism 9 (GraphPad Software, San Diego, CA). All *P* values, SEMs and *t*-statistics are shown on graphs. Welch’s unpaired one-tailed *t* test was used to compare means between the two groups (Tg-AD males and Tg-AD females) for IHC (Fig. 2, 4, 5) and WB (Fig. 6). Ordinary one-way ANOVA with Tukey’s post-hoc analysis was used to compare plaque burden in the four hippocampal regions (Fig. 2 C and D). Multi-factor comparisons for aPAT (Fig. 1) and microglia (Fig. 3) were performed using a two-way repeated measure analysis of variance (ANOVA), followed by a post hoc (Tukey’s) to access differences across individual training trials (FIG. 1) or the four groups of rats (Fig. 3). For image quantification, normalization of pixel intensity values across images was done utilizing the rolling ball algorithm [49].

**Figure 1.**
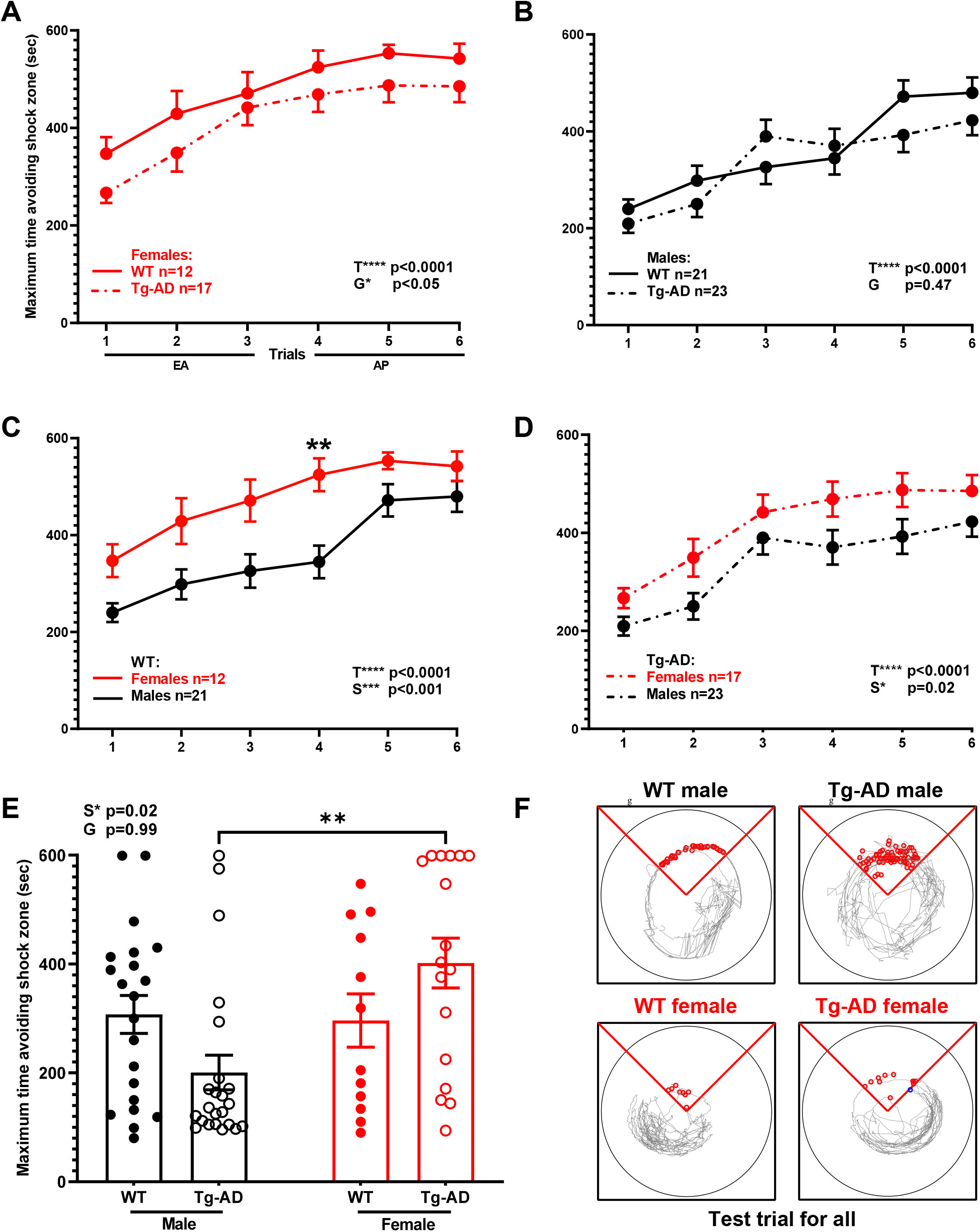
Females outperform males on spatial learning and memory assessed with the active place avoidance task (aPAT). Maximum time to avoid shock zone during training (A-D). There was an overall difference between WT and TG females (A) but not between WT and TG males (B). There was no difference in early acquisition (EA) and asymptotic performance (AP) between WT and TG in both sexes. A significant sex effect was observed as WT females performed better than WT males in EA (C), and TG females outperform TG males (D). The test trial 24 hours after training shows overall sex effect as TG females outperform TG males in maximum time to avoid shock zone (E). Track tracing of individual rat performances for the test trial across all groups (F). Repeated measures two-way ANOVA with Tukey’s post-hoc tests were used in 1A through 1D. Ordinary two-way ANOVA with Tukey’s post-hoc analysis was used in 1F. Females WT (n = 12), Tg-AD (n = 17); males WT (n = 21), Tg-AD (n = 23). *\*\* P* < 0.01. EA - Early Acquisition; AP – Asymptotic Performance. Overall statistical effects across trials (T*), sex (*S), or genotype (G*) are denoted.

**Figure 2.**
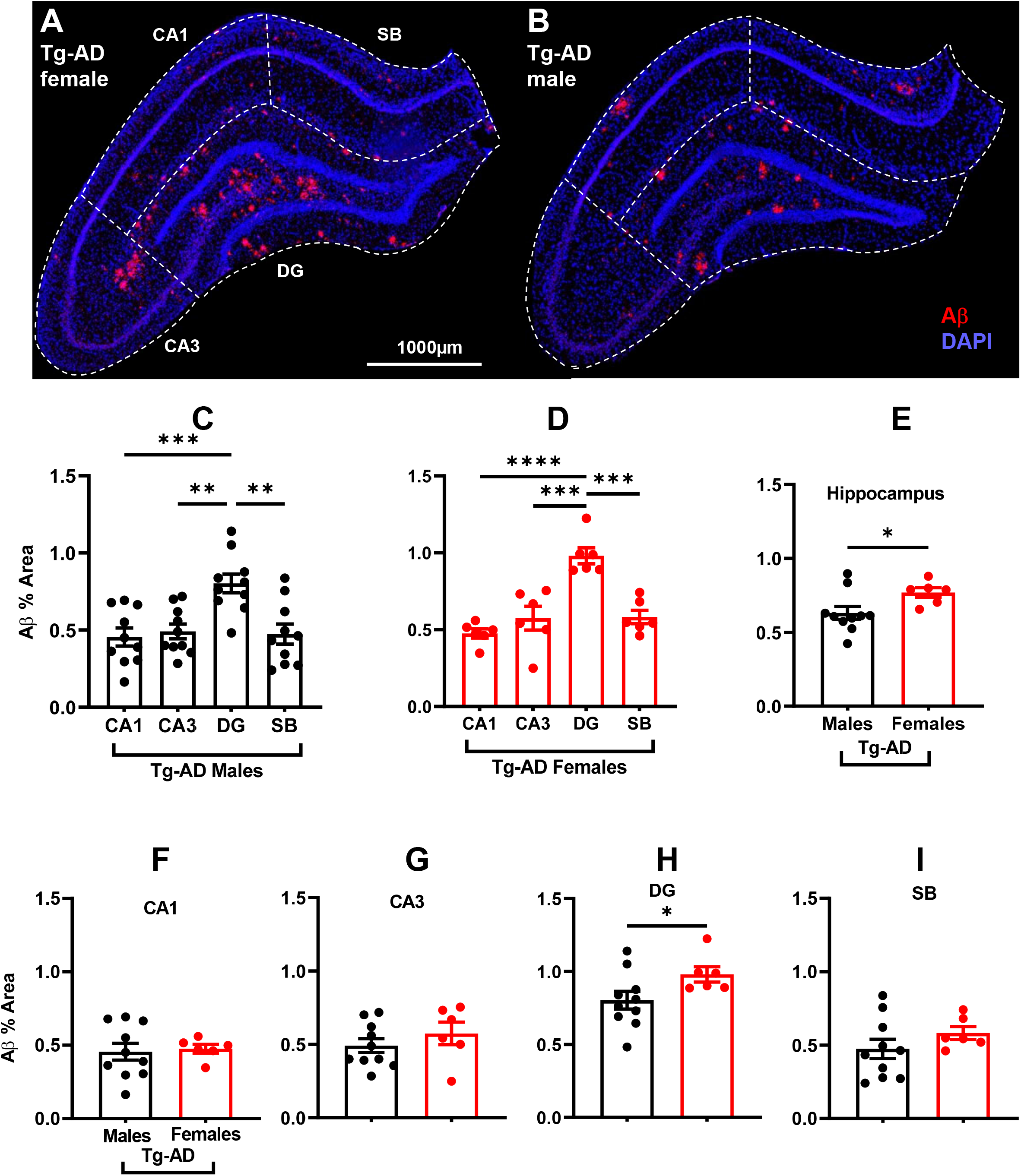
Aβ plaque burden is significantly higher in transgenic females than in males in total hippocampus and dentate gyrus (DG). Representative Aβ (red) and DAPI (blue) staining images for a transgenic female (A) and a transgenic male rat (B). Scale bar, 1000 µm. Hippocampal region perimeters are depicted in each image by white dashed lines. Plaque burden in transgenic males (C) and females (D) was significantly higher (∼1.7 fold) in DG than in other hippocampal regions. Significant increased plaque burden (*P < 0.05) was observed in TG females in whole hippocampus (E) and DG (H). No difference in plaque burden was observed in CA1, CA3, and SB (F-G, I). Ordinary one-way ANOVA with Tukey’s post-hoc analysis was used in 1C and D. Unpaired two-tail t-tests with Welch’s corrections were used in 1E through 1I. * *P* < 0.05. Tg-AD males (n = 10), Tg-AD females (n = 6). CA – Cornu Ammonis; DG – Dentate Gyrus; SB – subiculum.

**Figure 3.**
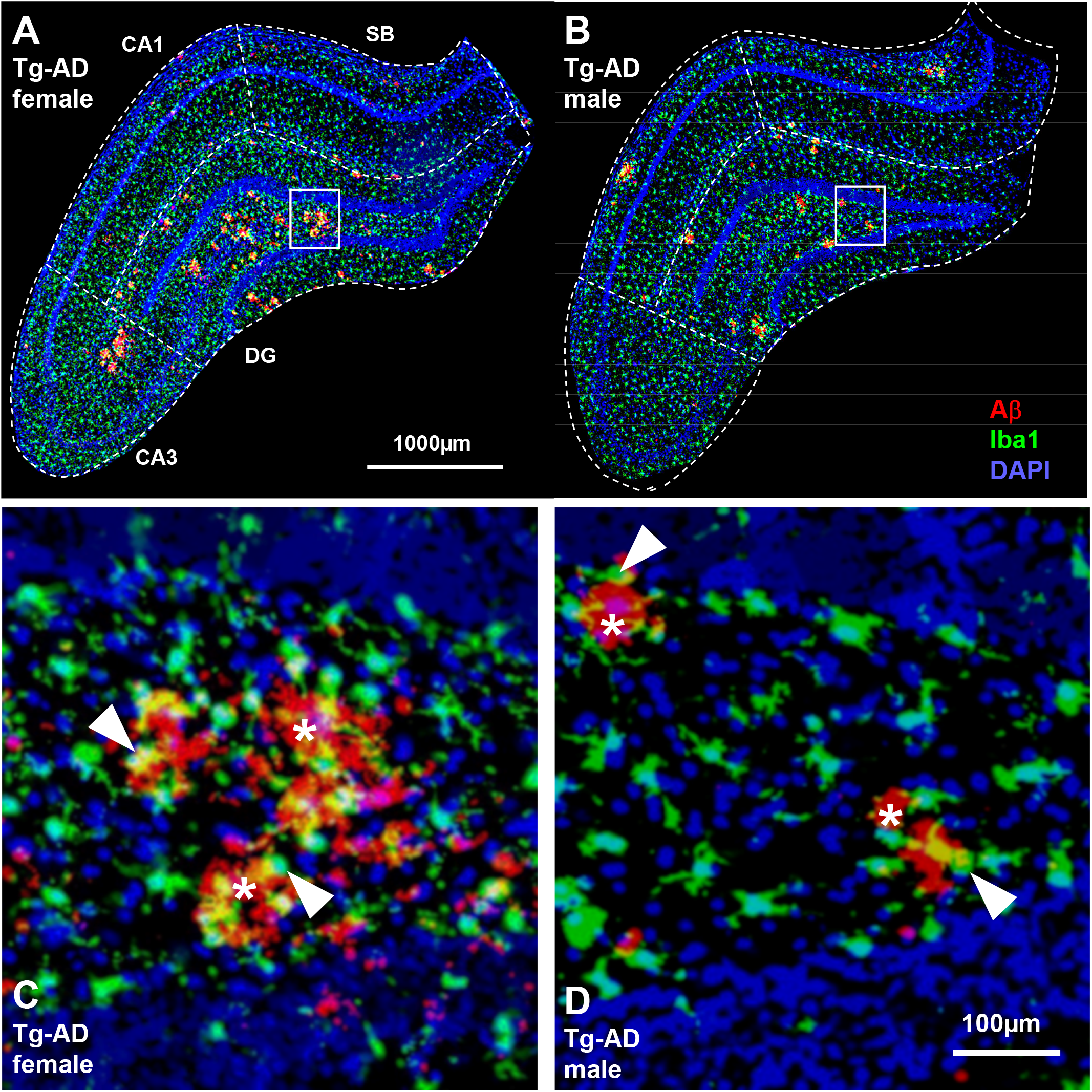
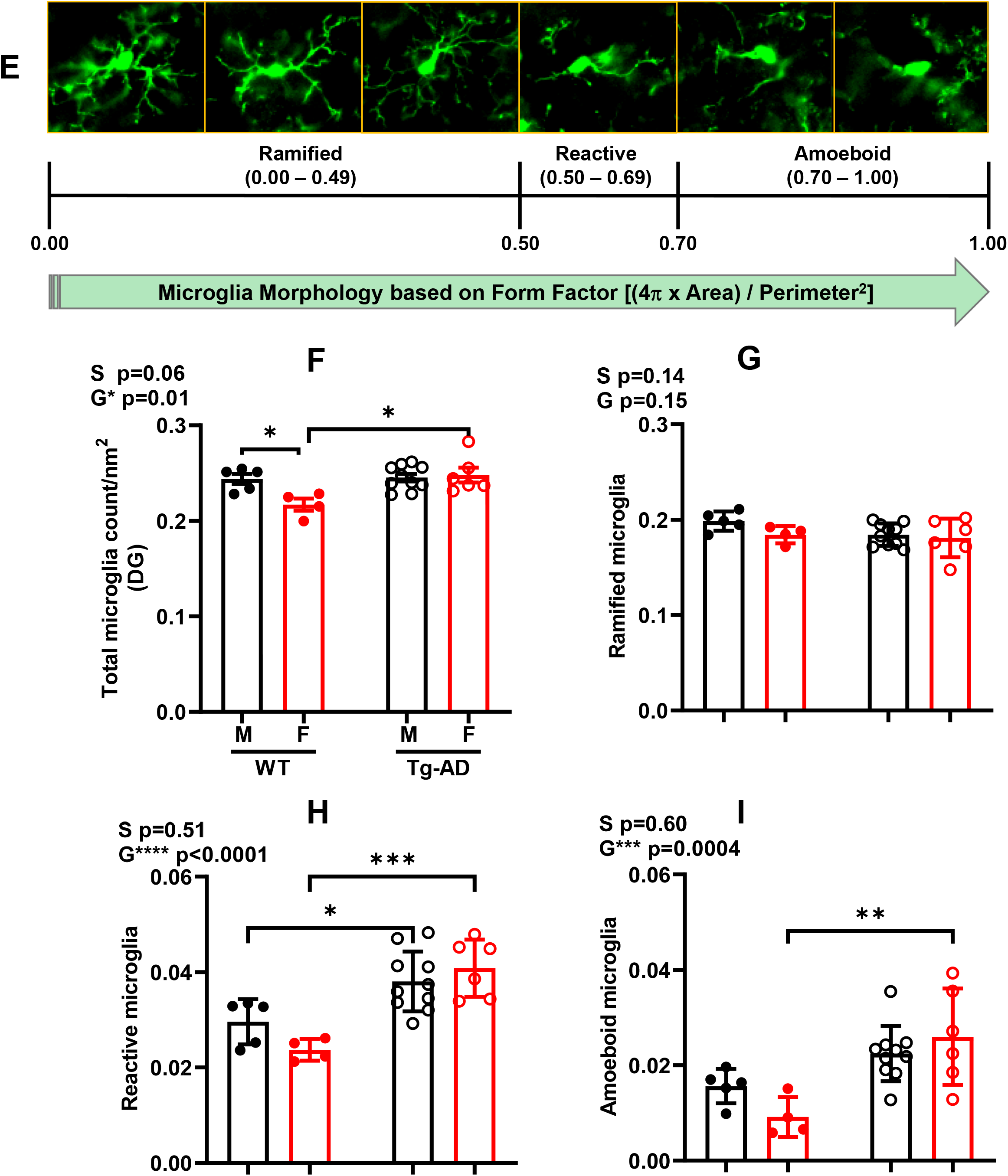
WT female rats have the lowest levels of microglia in the hippocampus compared with the three other groups, Tg-AD females as well as Tg-AD and WT males. IHC analysis for Aβ (red), microglia (green), and nuclei (DAPI, blue) of the right dorsal hippocampus of Tg-AD females (A) and Tg-AD males (B), 10x magnification, 1000 μm scale bar. Hippocampal region perimeters are depicted in each image by white dashed lines. Bottom panels (C and D) represent the magnification of the respective small white boxes depicted in A and B, 100 μm scale bar. Black asterisks show the Aβ plaques (red), surrounded by some microglia [white arrows show co-localization (yellow) of microglia and Aβ]. E, The three microglia phenotypes (ramified, reactive and amoeboid, all Iba1+) based on circularity (form factor) as explained under material and methods. WT females have significantly fewer microglia (counts/nm^2^) in the whole hippocampus (right dorsal) than the other three groups of rats (F). The number of ramified microglia was not significantly different in the four groups of rats (G). The number of reactive (H) microglia was significant lower in WT females and males compared to Tg-AD females and males. The number of amoeboid (I) microglia for WT females was significant lower compared to Tg-AD females and males. Ordinary two-way ANOVA with Tukey’s post-hoc tests were used in 3F through 3I. * *P* < 0.05, \*\**P* < 0.01, *** *P* < 0.001. Females WT (n = 4), Tg-AD (n = 6); males WT (n = 5), Tg-AD (n = 10). CA – Cornu Ammonis; DG – Dentate Gyrus; SB – subiculum.

**Figure 4.**
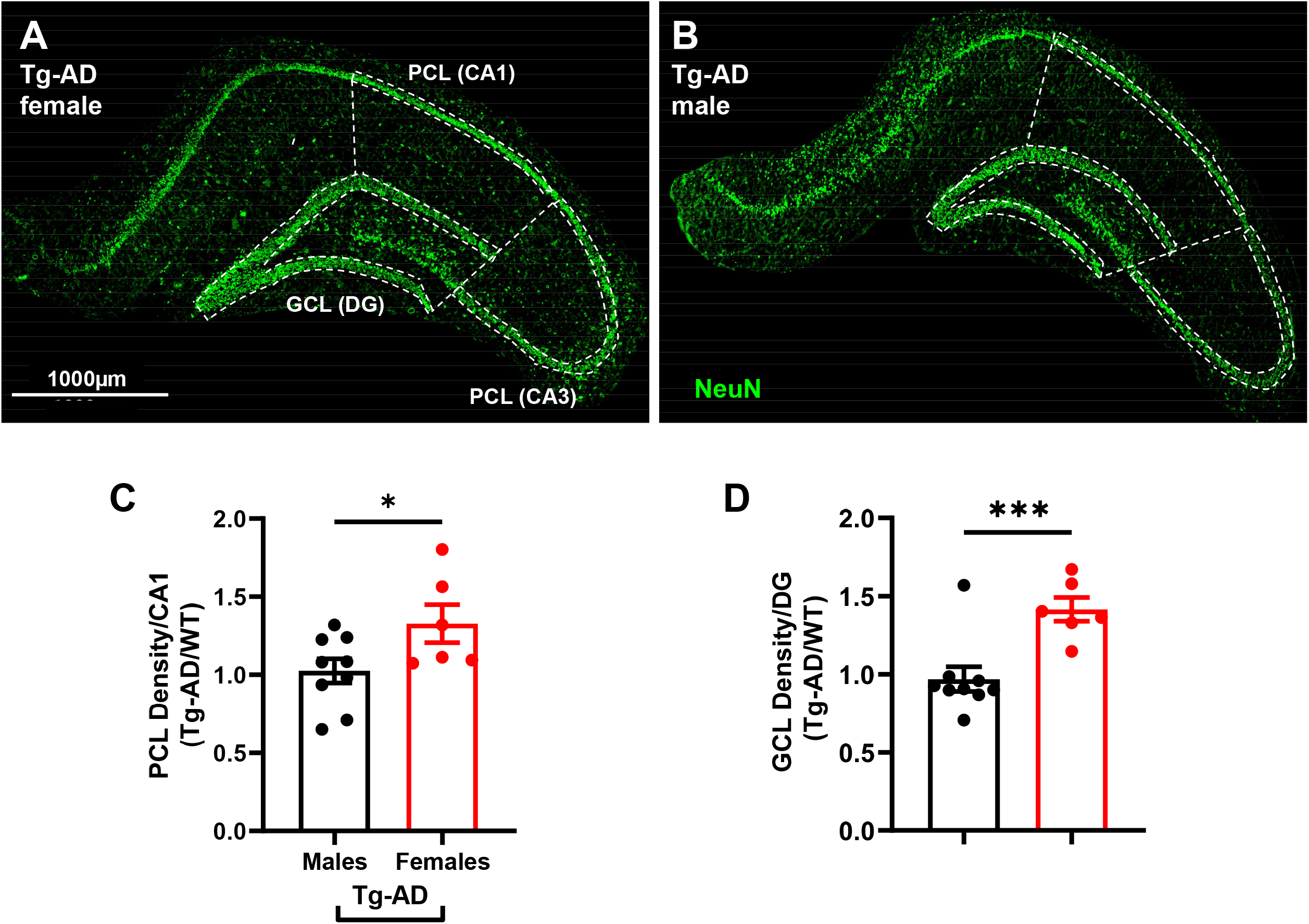
Tg-AD females have a higher hippocampal neuronal density than Tg-AD males. IHC analysis for mature neurons (green) of the right dorsal hippocampus of Tg-AD females (A) and Tg-AD males (B), 10x magnification, 1000 μm scale bar. Hippocampal pyramidal (PCL) and granular (GCL) neuronal cell layers are highlighted with white dashed lines. Tg-AD female rats had significantly higher neuronal density (NeuN staining) in the (C) PCL (CA1) and (D) GCL (DG) hippocampal regions compared to Tg-AD males. Unpaired one-tail t-tests with Welch’s corrections were used for comparison. * *P* < 0.05, *** *P* < 0.001. Tg-AD males (n = 9), Tg-AD females (n = 6). CA – Cornu Ammonis; DG – Dentate Gyrus.

**Figure 5.**
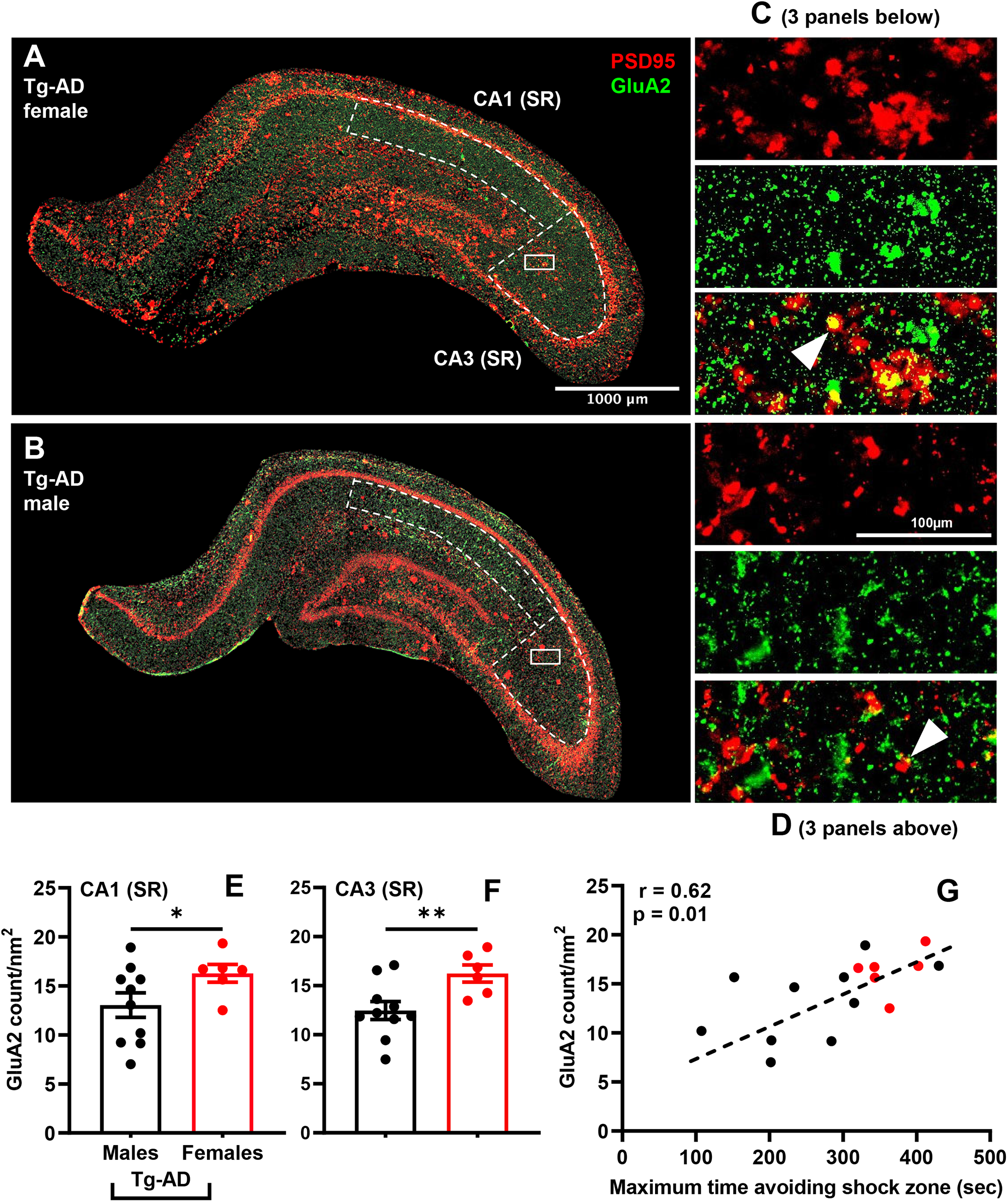
Tg-AD females exhibit increased levels of GluA2 in the hippocampus compared to Tg-AD males. IHC analysis for the AMPA receptor GluA2 subunit (green) and the scaffolding protein PSD95 (red) of the right dorsal hippocampus of Tg-AD females (A) and Tg-AD males (B), 10x magnification, 1000 μm scale bar. Smaller panels on the right (C and D) represent the magnification of the respective small white boxes depicted in A and B, 100 μm scale bar, showing the two stains (GluA2, green and PSD95, red) separately and co-localized (yellow, white arrows). Hippocampal CA1 SR and CA3 SR subregions are highlighted with white dashed lines. Tg-AD female rats had significantly higher GluA2 levels in the (E) CA1 (SR) and (F) CA3 (SR) hippocampal subregions compared to Tg-AD males. Unpaired one-tail t-tests with Welch’s corrections were used for the comparison. * *P* < 0.05, ** *P* < 0.01. (G) GluA2 levels (y-axis) directly correlates with cognitive behavior (x-axis). The correlation was evaluated by linear regression calculating the Pearson correlation coefficient. Tg-AD males (n = 9), Tg-AD females (n = 6). CA – Cornu Ammonis; SR - stratum radiatum.

**Figure 6.**
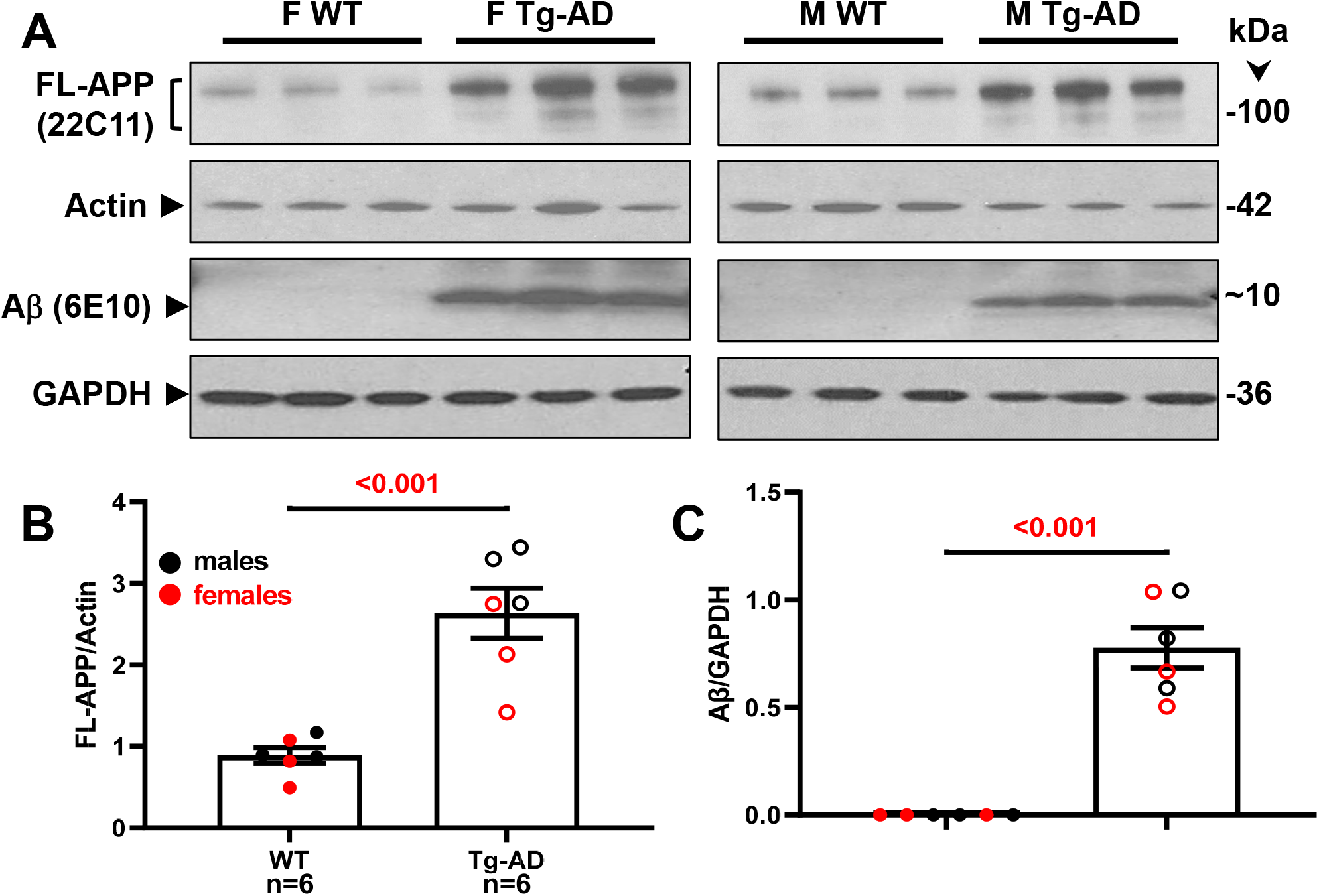
Tg-AD rats show enhanced FL-APP and Aβ peptide levels in the hippocampus at 9-months of age. (A) FL-APP (top panels) and Aβ levels (third panels from the top) were assessed by western blot analysis in whole left hippocampal (combined ventral and dorsal) homogenates from 9-month WT and Tg-AD female (F, 3 of each genotype) and male (M, 3 of each genotype) rats. Actin (second panels from the top) and GAPDH (bottom panels) detection served as the respective loading controls. FL-APP (B) and Aβ (C) levels were semi-quantified by densitometry. Data represent the percentage of the pixel ratio for FL-APP and Aβ over the respective loading controls for Tg-AD compared to WT (value of one). Values are means ± SEM from 6 rats per genotype (males and females combined). Significance (*p* values shown on graphs) estimated by an unpaired one-tail t-tests with Welch’s corrections. There was no significant difference between males and females for both genotypes.

## Results

### Female Tg-AD and WT rats outperform male counterparts at 9-months of age in spatial learning and memory

We performed aPAT on 9-month old WT and Tg-AD male and female rats in order to assess their spatial memory retention at this early stage of AD pathology. Animals were habituated to the testing apparatus and visual cues of the room, and then subjected to six training trials followed by a test trial with the shock off. The maximum time avoiding shock zone measure showed significant improvement across trials for the four groups of rats as they progressed through the trials with the best performance displayed on trials 5 and 6 (Fig. 1A-D).

For the maximum time avoiding shock zone during training, WT females performed better overall than Tg-AD females (Fig. 1A; *F* _(1,24)_ = 13.83, *P < 0*.*05*), there was no difference between the WT and Tg-AD female in early acquisition (EA) (*P = 0*.*32*) and asymptotic performance (AP) (*P = 0*.*51*). No differences were detected between WT and TG males across all 6 trials (overall) or in early acquisition (EA) and asymptotic performance (AP) [Fig. 1B, *F* _(1,24)_ = 0.53, *P* = 0.47; EA (*P = 0*.*99*); AP (*P = 0*.*71*)]. We also found that WT and Tg-AD female scores were superior to WT and Tg-AD males respectively for the maximum time to avoid shock measurement (Fig. 1C; *F* _(1,31)_ = 17.87, *P < 0*.*001*; with post-hoc differences at trial 4 (*t* = 179.9, *p* = 0.005) and Fig. 1D; *F* _(1,31)_ = 6.08, *P = 0*.*02*). This sex-dependent tendency was replicated in test trials 24 hours after training, with Tg-AD females performing better than Tg-AD males (Fig. 1E, *t* = 201.1, *P = 0*.*002*). Figure 1F shows the track tracing of individual rat performances for the test trial across all groups.

### Tg-AD females exhibit greater Aβ plaque burden than Tg-AD males in whole hippocampus and DG

Tg-AD rats express human APP and thus are prone to AD pathology, notably the accumulation of Aβ peptides forming extracellular plaques [7]. Thus, we performed immunohistochemical analysis for Aβ in WT and Tg-AD rats. As expected, no plaques were detected in WT rats of both sexes (not shown). Significant plaque load was observed in both male and female Tg-AD rats showing that these rats exhibit amyloid pathology already at 9-months of age (Fig. 2A and B, amyloid plaques, red; DAPI, blue).

As the course of the disease from early to late stages is proposed to spread from the entorhinal cortex (layer II) → dentate gyrus → CA3 → CA1 [30], we compared plaque load in four hippocampal regions, CA1, CA3, DG, and SB. All hippocampal regions presented amyloid plaques, but in different amounts. One-way ANOVA followed by Tukey’s post hoc analysis showed a regional effect, in that amyloid plaque burden was at least 1.6 fold higher in the DG region than in any of the other hippocampal regions both in male (Fig. 2C) and female Tg-AD rats (Fig. 2 D). This regional amyloid plaque distribution observed in the Tg-AD rats at 9-months of age supports that the spread of pathology regarding amyloid plaques recapitulates early stage AD.

We also compared the sex differences in amyloid plaque pathology, as sex discrepancies were previously reported for male and female patients in the development of AD as well as for other rodent models of AD [6, 35]. Our results show a significant increase in the % area amyloid plaques in female Tg-AD rats compared to male littermates in the whole hippocampus (Fig. 2E) and in the hippocampal DG region (Fig. 2H). In the other hippocampal regions no sex-dependent differences were observed in amyloid plaque load (Fig. 2F, G and I).

### Tg-AD females show increased microgliosis in whole hippocampus and DG compared to WT females, complimentary to Aβ plaque burden

We performed an Iba1 immunohistochemical analysis for microgliosis across CA1, CA3, DG and SB in WT and Tg-AD male and female rats (Fig. 3A and B, whole hippocampus, C and D respective high magnification panels, shown for Tg-AD rats only; amyloid plaques, red; microglia, green; amyloid plaques/microglia co-localization, yellowish, indicated by white arrow heads).

We observed a significant regional effect on microglia counts/nm^2^ with WT females exhibiting the lowest microglia counts among the four rat groups in the DG (Fig. 3F). There was no significant genotype effect on microglia count between WT and Tg-AD males, and no regional effects (Fig. 3F for DG). There were also no sex differences in microglia counts analyzed by two-way ANOVA (Fig. 3F).

Microglia have a remarkable variety of morphologies associated with their specific functions, and can be divided into three forms according to their functions and cell body circularity: ramified, reactive and amoeboid [26] (Fig. 3E). Highly ramified microglia can change to an amoeboid form on pathological stimulation [15]. Ramified microglia, which are considered the “homeostatic” neuroprotective form [16], were the most abundant followed by reactive and then amoeboid. No significant differences were detected in ramified microglia counts/nm^2^ among the four groups of rats (Fig. 3G). Reactive microglia are considered to be the activated pro-inflammatory form of microglia [16]. There were no sex differences for reactive microglia counts/nm^2^, although Tg-AD rats of both sexes presented significantly more reactive microglia than WT rats (Fig. 3H), which is consistent with the detection of amyloid plaques in the Tg-AD rats. Finally, amoeboid microglia are considered to be the neurotoxic and overactive form [16]. Although the number of amoeboid microglia was similar in WT and Tg-AD male rats, WT females displayed significantly fewer amoeboid microglia than Tg-AD females (Fig. 3I). These findings are consistent with the increased microgliosis that is observed in AD patients, in part due to active microglia extending their processes into the plaque core and clustering around amyloid plaques as shown in Fig. 3C and D [31, 39, 52].

### Tg-AD males display increased neuronal loss in CA1 and DG compared to Tg-AD females

We assessed neuronal density across hippocampal regions CA1 and CA3 pyramidal cell layers, as well as DG granular cell layer with NeuN (green) to quantify mature neurons (Fig. 4A and B, shown only for Tg-AD rats of both sexes). Compared to Tg-AD females, Tg-AD males exhibited significant fewer neurons after normalizing the PCL from CA1 and GCL from DG for Tg-AD rats to their respective WT values (Fig. 4C and D). We detected a significant difference in neuronal density between Tg-AD males and females only in the CA1 pyramidal cell layer (22.7% less, *t* = 2.09, *p* = 0.033) and DG granular cell layer (31.5% less, *t* = 4.05, p = 0.0007) (Fig. 4C and D).

### Increased levels of GluA2 in the hippocampus of Tg-AD females complement their superior cognitive performance compared to Tg-AD males

The levels of the AMPA receptor GluA2 subunit at the synapses are important factors for spatial memory [36, 44], as is the trafficking of AMPA receptors regulated by the scaffolding protein PSD95 that is enriched at glutamatergic synapses [27]. We performed IHC analyses of the hippocampus for GluA2 and PSD95 (Fig. 5 A and B, whole hippocampus, C and D respective high magnifications panels, shown for Tg-AD rats only; PSD95, red; GluA2, green; PSD95/GluA2 co-localization, yellowish, indicated by white arrow heads). The levels of GluA2 are higher in Tg-AD females than in Tg-AD males, specifically at the CA1 SR subregion (Fig. 5E, *t* = 2.10, *p* = 0.037) and CA3 SR subregion (Fig. 5F, *t* = 2.99, *p* = 0.005) depicted by the dashed lines in Fig. 5A and B. It is clear that the higher levels of GluA2 in Tg-AD females than in Tg-AD males correlates with improved spatial learning, as shown in Fig. 5G. These findings support the notion that higher regional hippocampal GluA2 levels in Tg-AD females than in Tg-AD males contributes to spatial memory maintenance independently of their higher plaque load.

### Tg-AD rats show enhanced FL-APP and Aβ peptide levels in the hippocampus

Our studies reveal that at 9 months of age the Tg-AD rats of both sexes express FL-APP at a significant higher level than their WT counterparts (Fig. 6D, top panel). This trend was observed in males (n = 3 for each genotype) and females (n = 3 for each genotype), and the values were normalized for actin (Fig. 6E, p < 0.001). Full length APP (FL-APP) was detected with the mouse monoclonal antibody 22C11, which reacts with human and rat, as well as other species (manufacturer’s specifications).

We also assessed Aβ levels in the same samples of rat hippocampal tissue with the mouse monoclonal antibody 6E10, which has a 3-fold higher affinity for human APP and Aβ compared to the rat counterparts (manufacturer’s specifications). Aβ peptides were detected in male and female Tg-AD rats but not in the WT littermates, as shown in Fig. 6D [third panels labeled with Aβ (6E10)], and semi-quantified in Fig. 6F (p < 0.001). The presence of Aβ plaques in all 9-month Tg-AD rats was confirmed by IHC analysis with the mouse monoclonal antibody 4G8, as shown for one female and one male Tg-AD rat in Fig. 2A.

## Discussion

The TgF344-AD (Tg-AD) rat model for AD is a robust model for basic and translational AD research, as it recapitulates all aspects of AD pathology including its age-dependency [39]. These rats develop hippocampal and cerebral cortical amyloid plaques, neurofibrillary tangles, astrogliosis, microgliosis, neuronal loss, and cognitive impairment [7]. Herein we characterize the age-dependent sequence of AD pathology between sexes in this transgenic rat model at an early stage of pathology. This is important since AD disproportionately affects women, who compared to men, exhibit a faster progression of hippocampal atrophy and neuronal loss, ultimately leading to cognitive deficits and memory impairment [29].

We show that there is a significant sex-dependent amyloid plaque load in 9-month Tg-AD rats. There was a significant higher % area level of plaques in female compared to male Tg-AD rats. This is important because accumulation of Aβ plaques is thought to be one of the main driving forces of neurodegeneration in AD, as it could contribute to synaptic dysfunction [34], neurofibrillary tangles as well as neuronal loss causing impaired memory and cognitive dysfunction. Consistent with our findings, 9-month-old Tg-AD rats were shown to have plaques in the entorhinal cortex, hippocampus and cortical arterioles [25] as well as the DG and CA1 regions [48]. In addition, 9-month-old TgF344-AD rats showed doubling of tau phosphorylation at Ser202/Thr205 and a 1.5-fold increase in tau phosphorylation at Thr231 [4, 25]. At 6 months of age TgF344-AD rats had reduced basal synaptic transmission in entorhinal-hippocampal synapses and an increase in Aß oligomers, hyperphosphorylated tau and activated microglia and astrocytes [48] as well as reduced hippocampal axon density at 6 months of age [20].

A number of other studies using different AD animal models showed that females display more amyloid plaques at different ages [23] and higher levels of tau phosphorylation at late stages [6]. The amounts of Aß in female APP/PS1 mice were significantly higher than in the males, but unlike our findings corresponded to reduced memory test performance [29] It is possible that these findings may reflect that Aβ deposits might have a greater effect on cognition in these AD models.

The sex differences exhibited by the Tg-AD rats is consistent with the discrepancy exhibited by male and female patients in the development of AD, as well as neuronal loss in rodent models [5, 32]. A sex effect was previously reported as AD and other dementias disproportionately affect women [24]. This discrepancy is observed through a faster progression of hippocampal atrophy in females with AD compared to men [29]. There are also regional hippocampal differences in neuronal loss in AD [48, 49].

Sex differences in the Tg-AD rat model were also reported where basal synaptic transmission deficits at CA3-CA1 synapses were observed in 9 months males and not until 12 months in females [48]. These data support a neuroprotective role for estrogen in this rat model. However, other studies showed that spatial memory impairment in Tg-AD rats was evident in 6- to 7-month-old animals but no sex differences were observed [40].

As AD is a form of dementia characterized by memory loss, we looked for amyloid pathology in the various regions of the hippocampus in the Tg-AD rats. We detected the presence of amyloid plaques in the CA1, CA3, DG and SB regions in the rats at 9-months of age. There was statistically significant difference between males and females only in the DG hippocampal region with female having more plaques than male Tg-AD rats. These data suggest that at 9-months of age, the amyloid pathology is prominent in the DG region. Additional studies are necessary to further characterize the progression of amyloid plaque pathology throughout the hippocampus as the transgenic rats age.

AD is one of the most common causes of dementia, as loss of memory is among the first symptoms reported by AD patients since the disease affects both working memory and long-term memory [20]. This cognitive deficit is expressed by the Tg-AD rat model, as the transgenic rats displayed significant spatial memory impairment relative to WT animals [6]. We assessed cognitive ability of the Tg-AD rats via aPAT, to identify genotype-dependent spatial memory impairment. Latency to 1st entrance, is the most stringent and sensitive trial related to memory, as it is the first trial following habituation, in which the goal is to allow rats to learn and remember the visual cues surrounding the platform. The higher the value, the longer it took the rat to enter the shock zone for the first time, supporting a sound level of spatial memory and recognition. Similarly, higher values for latency to 2nd entrance and maximum time avoiding shock, indicate remembering the shock zone location, and avoiding it for a prolonged period of time. Improved performance in the aPAT assessment is indicative of sound memory, as hippocampal inactivation impairs performance in this task [4]. All rats showed improved performance in all three measures by the final training trial, supporting that there are no significant cognitive and memory deficits at 9-months of age in these rats. It is likely that at 9-months of age, the limited AD pathology detected in the hippocampus of these transgenic rats is not sufficient to impair cognition. The behavioral deficits will most likely develop at a later age, when pathology is more advanced.

Previous findings in Tg-AD rats reported neurocognitive impairments in 5-month rats (pre pathology) by means of delayed non-match-to-sample task [35]. Although we observed poorer performance in Tg-AD compared to WT littermates, the difference was not significant. This result suggests that 9-months of age may be too early to observe significant cognitive decline via aPAT using this Tg-AD rat model. Neurocognitive impairment for this rat model was detected at 6-month age using a Barnes maze [6]. However, the Barnes maze is stressful on its own to the rats causing an increase in plasma corticosterone, thus causing test-induced responses that, by themselves, can affect cognition [18] adding an additional stress factor to the AD pathology.

In our studies, female WT and Tg-AD rats scored better with the aPAT assessment than their male littermates. In a previous study, Tg-AD rats showed spatial memory impairment at 4 to 5 months of age using Active Allothetic Place Avoidance (AAPA), which increased at 6- to 7 months [40]. However, unlike our studies, no sex effect was observed. Females of both genotypes exhibited superior latency to 1st entrance, latency to 2nd entrance, and maximum time to avoid shock than males. These aPAT results are in contrast with the immunohistochemical analysis, showing higher amyloid plaque load and gliosis in transgenic female rats than in males. This suggests that the progression of cognitive impairment is slower in females presenting itself later in age than in males.

AD is also associated with chronic neuroinflammation caused by overactive microglia [10]. The state of microgliosis in the Tg-AD rats at 9-months of age was determined. WT and Tg-AD rats displayed increased microglia count in the hippocampus. Similar to plaque load, there was a significant genotype effect on microglia numbers and % area between WT and Tg-AD females. Microglia are recruited to amyloid plaques for phagocytosis [3]. Microglia exhibit a remarkable variety of morphologies that can be associated with their particular functions [22]. Since gliosis is implicated in the pathology of AD, we hypothesized that microglia morphology may be altered. An examination of individual microglia silhouettes revealed that Tg-AD rats of both sexes had more reactive microglia than WT but no sex differences. These findings are consistent with the increased levels of amyloid plaques in Tg-AD rats. Interestingly, the levels of amoeboid microglia which are considered to be the neurotoxic ones, was significantly greater in Tg-AD female than WT females. These results support findings that AD patients show increased gliosis.

Upon recognition of a threat to the CNS, such as amyloid plaques, microglia are recruited to the site of damage and extend their processes into the plaque core, associating with amyloid plaques to perform their phagocytic activity and clear the plaques [27, 37, 50]. The extensive gliosis detected in the Tg-AD females may cause higher levels of neuroinflammation, leading to neuronal loss and further exacerbating the AD pathology in females [19, 31, 42]. Thus, higher levels of neuroinflammation could be another factor contributing to the disproportionate effect of AD on women, and are manifested as the higher plaques and microglia that we observe in 9-month female Tg-AD rats as an indication of their sensitivity to the disease. In contrast, female Tg-AD rats had more neurons than Tg-AD males which may explain the greater cognitive performance observed in females.

In this study, we characterized the AD-like pathology developed by Tg-AD rats at 9-months of age. Overall, we established that transgenic female rats exhibited higher levels of amyloid plaques and gliosis, thus exacerbated AD-like pathology compared to males. In contrast, females performed better than males in a hippocampal-dependent cognitive task. Estrogen is well known for being neuroprotective but has a complicated relationship when it comes to AD [51]. In this regard, estrogen was shown to improve spatial learning in APP/PS1 mice without affecting the accumulation of beta amyloid plaques [22]. Nevertheless, estrogen was shown to protect against apoptosis induced by Aβ in hippocampal neurons [37]. A relationship between estrogen and increased GluA2 was demonstrated in gonadectomized rats where estrogen treatments resulted in a two-fold higher increase in hypothalamic GluA2/3 expression in females compared to males [12]. The ability of estrogen to enhance learning and memory was further shown by the demonstration that estrogen stimulated the trafficking of GluA2 to mushroom spines via the PI3 signaling pathway [2].

Our studies show that the improved cognition in 9-month Tg-AD females was paralleled by increased levels of GluA2. GluA2 plays an important role in maintaining synaptic plasticity and estrogen can regulate its levels. This notion is supported by studies showing that a reduction in synapse dysfunction and impaired cognition in 3xTg-AD was accompanied by an increase in the levels of GluA2 [41]. Other studies with male and female rats showed that a significant correlation existed between retention tests scores and synaptic GluA2 levels in the hippocampus [43]. Similarly, an improvement in spatial memory in the hippocampus in Tg 2576 mice was associated with increased expression of both GluA1 and GluA2 at synapses without affecting Aβ protein levels [17]. Also, mitigation of memory deficits resulted in increased GluA1 and GluA2 levels in the 3xTg-AD mouse model, suggesting that AMPA receptors play a critical role in maintaining synaptic plasticity and memory [41]. While these finding support a critical role for GluA2 in synaptic plasticity, GluA2 is significantly expressed in the human post-mortem hippocampus of AD patients relative to controls, specifically in the stratum moleculare of the DG [54].

In conclusion, our results, suggest that estrogen protects female Tg-AD rats against cognitive dysfunction at early stages of AD but has little effect on other pathological deficits. This estrogen-mediated protection is associated with an increase in GluA2, suggesting that increased activity of AMPA receptors may preserve synaptic plasticity at the onset of AD. This hypothesis needs to be further investigated in future studies.

## Acknowledgments

This work was supported in part by NIH/NIA R01AG057555 to LX, NIH training grant R25GM060665 to support OC and KN, and the City University of New York (Ph.D. program in Neuroscience, Graduate Center). We thank the technical support of Charles Wallace and Giovanni Oliveros (both PhD candidates) in the Department of Biological Sciences at Hunter College, CUNY.

## Abbreviations

Aβ: amyloid-beta
AD: Alzheimer’s disease
APP: amyloid precursor protein
BACE1: β-secretase 1
CA: cornu ammonis
DG: dentate gyrus
GAPDH: glyceraldehyde 3-phosphate dehydrogenase
GCL: granular cell layer
HL: hilar
Iba1: ionized calcium binding adaptor molecule 1
NeuN: neuronal nuclei, neuronal marker
PCL: pyramidal cell layer
SB: subiculum
SEM: standard error of the mean
SR: stratum radiatum
TgF344-AD = Tg-AD: transgenic rat model of Alzheimer’s disease
WT: wild type

## Author Contributions

OC, MFP, PR, and PS conceived the project and designed the experiments; OC performed most of the experiments and analyzed the data; KN contributed to the behavioral assays, western blot and data analyses; OC, MFP, PR, and PS wrote the manuscript. All authors approved the manuscript for submission.

## Data Availability

Data supporting the findings of this manuscript are available from the corresponding authors upon request

